# From Adalimumab to orally administered secretory IgA: preclinical characterization of ABC-301

**DOI:** 10.64898/2026.11.16.766909

**Authors:** M. Brousse, A. Pintat, B. Lejeune, O. Favre-Bulle, JM. Herbert, A. Cuvillier

**Author notes:** **CONTACTS INFORMATION: JM. Herbert**, ABCELY, 2, rue Crucy, 44000 Nantes,**, A. Cuvillier,** ABCELY, 2, rue Crucy, 44000 Nantes. JMH and AC contributed equally to this work.

## Abstract

Reformatting therapeutic immunoglobulin G (IgG) antibodies into secretory immunoglobulin A (sIgA) may provide an alternative architecture for mucosal drug delivery. We characterized ABC-301, a recombinant human sIgA1 incorporating the antigen-binding regions of Adalimumab, and evaluated its human tumor necrosis factor alpha (TNFα) neutralization, oral activity in experimental colitis, tissue biodistribution, and susceptibility to patient-derived anti-Adalimumab antibodies. ABC-301 retained human TNFα binding and functional neutralization *in vitro*. Its whole-molecule molar neutralization IC_50_ was approximately 3.3-fold lower than that of Adalimumab, a difference of approximately 1.6-fold after normalization to theoretical binding-site number. In dextran sulfate sodium (DSS)-induced colitis, oral ABC-301 attenuated colon shortening and reduced colonic inflammatory myeloid populations and tissue inflammatory biomarkers relative to untreated DSS controls. Following oral administration, iodine-125-associated radioactivity was predominantly detected in gastrointestinal compartments. In seven patient-derived anti-Adalimumab antibody preparations, ABC-301 showed reduced functional susceptibility to inhibition compared with Adalimumab. These findings provide a preclinical proof of concept for IgG-to-sIgA reformatting.

## INTRODUCTION

Inflammatory bowel diseases (IBD), principally Crohn’s disease (CD) and ulcerative colitis (UC), are chronic relapsing inflammatory disorders of the gastrointestinal tract. Historically recognized as distinct clinicopathological entities, CD is characterized by discontinuous, frequently transmural inflammation that can affect the entire gastrointestinal tract, whereas UC predominantly involves continuous mucosal inflammation of the colon. Despite these anatomical and immunopathological differences, both diseases are characterized by disruption of epithelial barrier integrity and dysregulated activation of innate and adaptive immune responses (1,2).

TNF-alpha is a central inflammatory mediator in IBD and promotes leukocyte recruitment, cytokine amplification, epithelial dysfunction, and tissue injury (3) Therapeutic antibodies targeting TNF-alphfa have transformed the management of moderate-to-severe IBD (4,5). However, a substantial proportion of patients show primary non-response or subsequently lose response. Anti-drug antibodies (6) pharmacokinetic variability (7) and TNF-independent inflammatory pathways can contribute to treatment failure (8). In addition, currently used anti-TNFα biologics are delivered systemically, despite the intestine being the principal site of disease activity.

Secretory IgA is the predominant immunoglobulin class at mucosal surfaces. Its polymeric architecture, association with the J chain and secretory component, and relative resistance to proteolysis are adapted to mucosal environments. These properties raise the possibility that a therapeutic sIgA could be administered orally and preferentially expose gastrointestinal tissues to an antibody while limiting systemic exposure.

ABC-301 is a recombinant human anti-TNF-alpha sIgA1 generated by transferring the variable regions of heavy and light chains of Adalimumab into a human IgA1-based secretory antibody format. The molecule consists of two IgA units associated with a J chain and secretory component. This design tests a broader pharmacological principle: an established therapeutic IgG paratope can potentially be preserved while antibody isotype and route of administration are changed to favor mucosal exposure. Such a strategy could be relevant beyond TNFα for therapeutic IgGs whose systemic administration is associated with limited delivery to a mucosal site of action, systemic exposure, or other format-related constraints.

Following systemic administration, the exposure of inflamed intestinal tissues to Adalimumab is governed by multiple pharmacokinetic processes, including absorption after subcutaneous administration, systemic distribution, inflammation-associated clearance (9) and tissue penetration. Several studies in patients with IBD have revealed heterogeneous serum-to-intestinal drug distribution, depending on the stage of diseases due to “sink effect” in inflamed tissues for anti-TNFα drug (10). Despite its established clinical efficacy in inflammatory bowel disease, Adalimumab treatment is frequently complicated by the development of anti-drug antibodies (ADAs), which are associated with reduced circulating drug concentrations, loss of therapeutic response and treatment discontinuation. Prospective studies have reported anti-Adalimumab antibody development in approximately 20–35% of patients, with antibodies detectable as early as two weeks after treatment initiation. Importantly, ADA positivity has been associated with a threefold increased risk of Adalimumab discontinuation due to loss of response, highlighting immunogenicity as a clinically relevant limitation of long-term anti-TNFα therapy (6,11–13). Although ABC-301 is engineered based on the same anti-TNFα paratope as Adalimumab, its distinct sIgA1 architecture may alter the recognition and functional inhibition by anti-Adalimumab antibodies.

Here, we report the preclinical characterization of orally administered ABC-301, including TNF-alpha binding and neutralization, efficacy in DSS-induced colitis in human CD89-transgenic mice, local inflammatory biomarkers, ADA responses in mice, and sIgA reformatting benefit toward anti-ADA neutralization and gastrointestinal biodistribution. ABC-301 represents the first successful validation of this IgG-to-sIgA reformatting platform, demonstrating that a clinically validated IgG paratope can be converted into a functionally orally active secretory IgA. We investigated whether IgG-to-sIgA1 reformatting could reduce susceptibility to ADA-mediated neutralization, potentially offering a therapeutic advantage in patients who develop resistance to Adalimumab. This proof of concept supports the applicability of an adaptable platform, in which each sIgA candidate can be specifically optimized and validated according to the properties of the parental antibody and its therapeutic application.

## MATERIAL AND METHODS

### ABC-301 design and production

ABC-301 is a recombinant human secretory IgA1-kappa antibody containing the anti-TNF-alpha complementarity-determining regions derived from Adalimumab. Heavy chain, light chain, joining chain, and secretory component were co-expressed in suspension-adapted CHO cells. Research-grade material was purified using Protein L affinity chromatography followed by preparative size-exclusion chromatography. The structural integrity of ABC-301 was assessed by SDS-PAGE using NuPAGE™ 3–8% Tris-Acetate gels (Invitrogen) according to the manufacturer’s instructions. Proteins were visualized using Colloidal Blue staining, and molecular weights were estimated using HiMark™ prestained protein standards (Invitrogen).

### TNF-alpha binding assay

High-binding 96-well ELISA plates (MaxiSorp®, Thermo Fisher Scientific) were coated overnight at 4°C with recombinant human TNFα (0.75 µg/mL) diluted in phosphate-buffered saline (PBS). After washing, nonspecific binding sites were blocked with 3% bovine serum albumin (BSA) in PBS. Purified ABC-301 or Adalimumab were added to the wells and incubated for 1 hour at 37°C. Following washing, bound antibodies were detected using horseradish peroxidase- or alkaline phosphatase-conjugated anti-human IgA or anti-human IgG secondary antibodies. After substrate enzymatic cleavage, absorbance was measured at 450 nm. Samples were considered positive when the absorbance exceeded the predefined cutoff established from negative-control wells. Half-maximal effective concentrations (EC50) were derived from concentration-response curves.

### L929 TNF-alpha neutralization assay

Functional TNF-alpha neutralization was evaluated using TNF-alpha-sensitive L929 fibroblast-like cells. TNF-alpha induces apoptosis in this cell line; inhibition of cell death therefore provides a measure of neutralizing activity. ABC-301 monomer-enriched and oligomer-enriched fractions were compared with Adalimumab, and apoptosis inhibition was quantified by flow cytometry. Half-maximal inhibitory concentrations (IC_50_) were calculated from concentration-response curves.

### ADA neutralization test

Anti-Adalimumab antibody preparations from seven Adalimumab-treated patients were investigated using the L929 TNFα-dependent cell-death assay. TNFα was used at 7.5 ng/mL, and the therapeutic antibody (Adalimumab or ABC-301) concentration was stated as 2 × 10⁻¹⁰ mol/L while ADA concentrations were varied over nominal therapeutic antibody/ADA molar ratios ranging from 0.5:1 to 19,200:1. For each patient preparation and therapeutic antibody, the ratio at which ADA-mediated functional inhibition was no longer detectable was recorded. The comparison was paired within each patient.

### Animals, DSS colitis, and ethics

Experimental colitis was induced in male and female C57BL/6 mice transgenic for human CD89 by administration of 1.5% dextran sulfate sodium (DSS) in drinking water ad libitum for five days. Animals were assigned to healthy, untreated DSS, ABC-301-treated DSS, or Adalimumab-treated DSS groups. ABC-301 was administered by oral gavage once daily from day 6 through day 12 at 2, 1, 0.5, or 0.2 mg/kg. Adalimumab was administered subcutaneously at 2 mg/kg on day 6. Body weight and blood in stools were monitored daily. Animals were euthanized on day 13; blood, colon, fecal material, and spleen were collected. Colon length was measured after removal of fecal contents and rinsing with 0.9% NaCl.

### Flow cytometry

Single-cell suspensions were prepared from emptied and washed colon tissue by mechanical dissociation. Cell suspensions were filtered through 40-µm cell strainers, and erythrocytes were lysed. Cells were subsequently washed and resuspended in flow cytometry staining buffer. After cell counting, nonspecific Fc receptor binding was blocked using an Fc receptor–blocking reagent. Cells were then stained with a viability dye together with fluorochrome-conjugated monoclonal antibodies specific for myeloid lineage markers. Flow cytometric analyses were performed on viable singlet cells following sequential exclusion of debris, doublets, and dead cells. Neutrophils were identified as CD11c^-^CD11b^High^Ly6G^+^ cells, inflammatory monocytes as CD11b^+^Ly6G^-^F4/80^-^ cells, macrophages as CD11c^High^CD11b^int^F4/80^+^Ly6G^-^ cells and M1 macrophages as CD11c^+^CD11b^+^F4/80^+^CD80^+^. Fluorescence-minus-one (FMO) controls, together with unstained and single-stained compensation controls, were used to establish gating strategies and fluorescence compensation. Data were acquired using a CytoFLEX (Beckman Coulter) and analyzed with BD FACSDiva™ Software (BD Biosciences). Cell populations were expressed as the percentage of viable leukocytes recovered from organ.

### Calprotectin and Lipocalin-2 quantification

Calprotectin and lipocalin-2 were quantified in intestinal tissue and feces, by ELISA kits according to manufacturer’s instructions. (Calprotectin: DuoSet™ ELISA Development System Mouse S100A8/S100A9 Heterodimer, R&D Systems, Catalog Number: DY8596-05 and Lipocalin: Mouse Lipocalin-2/NGAL Immunoassay, R&D Systems, Catalog Number MLCN20).

### Murin Anti-drug antibody measurements

ADA responses were assessed by ELISA in plasma, intestinal tissue, and feces from DSS-treated animals receiving ABC-301 (7 oral day doses at 2, 1, 0.5 and 0.2mg/kg) or Adalimumab (1 sc dose at 2mg/kg). ABC-301 or Adalimumab was coated in High-binding 96-well ELISA plates (MaxiSorp®, Thermo Fisher Scientific) at 1µg/mL overnight in PBS 1X. Plasma, intestinal tissue and feces samples were incubated at 1/100 dilution for 1h at 37°c. Murin Ig were detected by anti-mouse Ig secondary antibody coupled to Alkaline Phosphatase (1/1000 dilution). After substrate enzymatic cleavage, absorbance was measured at 450 nm. Samples were considered positive when the absorbance exceeded the predefined cutoff established from negative-control wells.

### Radiolabelled biodistribution

ABC-301 was radiolabeled with iodine-125. In healthy mice, a single 40 microgram oral dose was administered by gavage and blood and tissues were collected at 2, 4, 8, 16, and 24 h. In a separate comparison in DSS-induced human CD89-transgenic mice, a 40-microgram oral dose was administered, and blood and tissues were collected at 2, 8, 16, 24, and 48 h. Tissue-associated radioactivity was quantified using a calibrated automated gamma counter (Hidex AMG). Gastrointestinal tissues, gastrointestinal washes, liver, spleen, and blood were analyzed for ABC-301-associated radioactivity, normalized to organ weight.

## Statistical analysis

Data are presented as mean ± standard deviation (SD) or as median [IQR]. Statistical analyses were performed using GraphPad Prism 10 (GraphPad Software, San Diego, CA, USA). Comparisons between two independent groups were performed using the two-tailed Mann–Whitney *U* test. Survival was summarized using Kaplan-Meier estimates. All tests were two-sided, and *P* values <0.05 were considered statistically significant. The paired patient-derived ADA comparison was reported as a paired t test on log-transformed therapeutic antibody/ADA thresholds, yielding a geometric mean paired ratio and 95% confidence interval. Statistical significance is indicated as follows: ns, not significant; *\* p* < 0.05; *\*\* p* < 0.01; *\*\*\* p* < 0.001 and *\*\*\*\* p* < 0.0001.

## RESULTS

### IgG-to-sIgA transformation fully preserves anti-TNFa avidity and functional TNFα-neutralizing activity

ABC-301 is a recombinant human secretory IgA1-kappa antibody containing the anti-TNF-alpha paratope derived from Adalimumab. Human Apha heavy chain, human Kappa light chain, human joining chain, and human secretory component were co-expressed in suspension-adapted CHO cells. Research-grade material was purified using Protein L affinity chromatography followed by preparative size-exclusion chromatography. The principal purified sIgA species consists of a monomeric form composed of two IgA units associated with one J chain and one secretory component as illustrated in the graphical representation (FIG 1A.) and the band at ∼350KDa identified on electrophoretic acrylamide gel (FIG 1B.). Two high-molecular-weight oligomeric forms are also purified in very minor amounts: a trimeric form consisting of three IgA units, one J chain and one secretory component, and a tetrameric form comprising four IgA units, one J chain and one secretory component. Finally, a band at approximately 150 kDa is characteristic of a very small residual amount of monomeric IgA that has not undergone dimerization or assembly into sIgA1.

**Figure 1.**
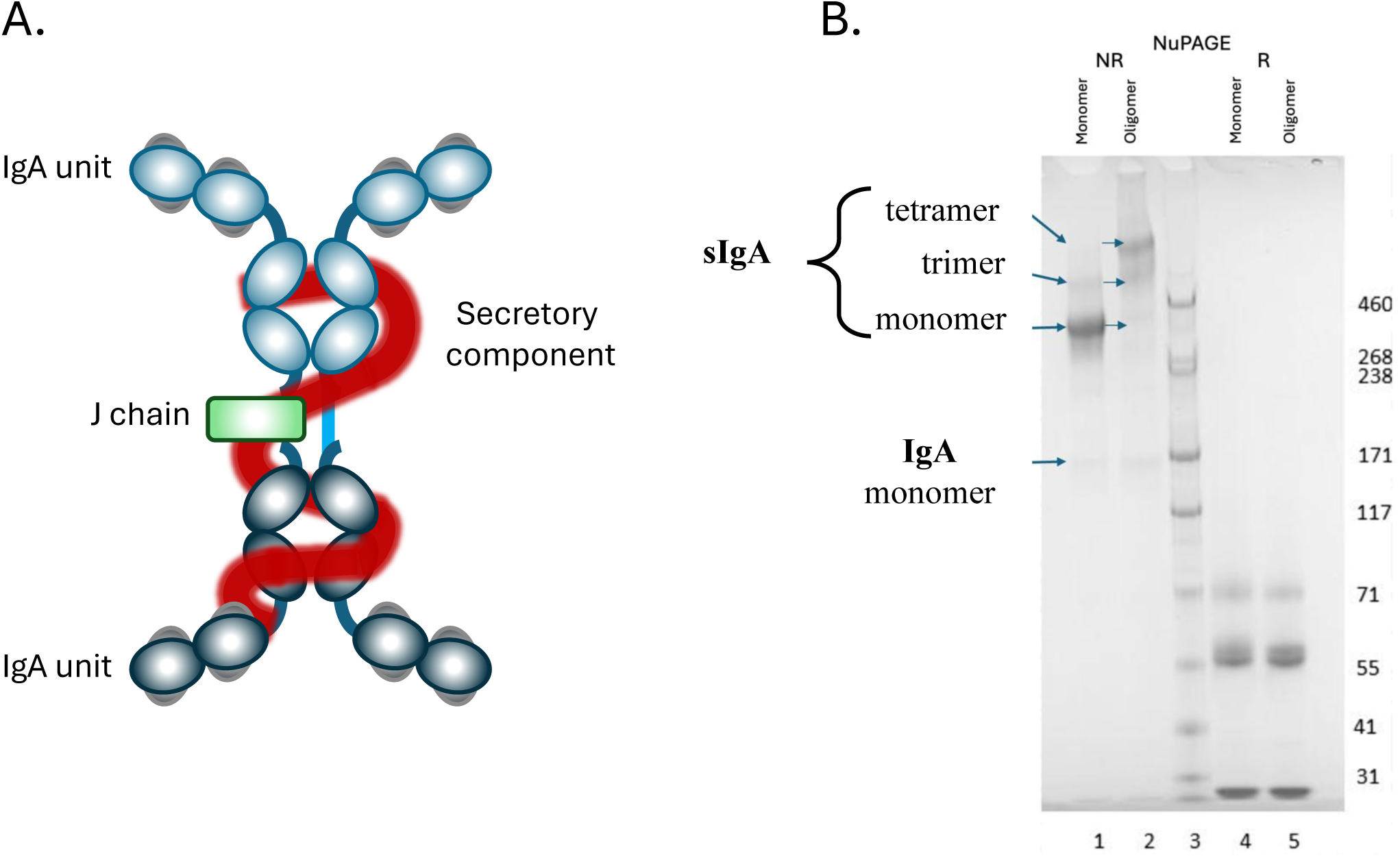
Molecular architecture and biochemical characterization of ABC-301, an adalimumab-derived secretory IgA. **(A)** Schematic representation of ABC-301, a recombinant human secretory IgA1 (sIgA1) incorporating the antigen-binding variable regions of Adalimumab. The predominant molecular species consists of two IgA units assembled through a joining (J) chain and associated with a secretory component (SC), resulting in a dimeric IgA assembly with four potential antigen-binding sites. **(B)** Electrophoretic characterization of purified ABC-301 preparations showing the predominant secretory IgA species, with an apparent molecular mass of approximately 350 kDa. Protein preparations were analyzed by SDS-PAGE using 3–8% Tris-acetate gels and visualized by colloidal blue staining. Molecular masses were estimated using protein molecular-weight standards.

Binding to human TNFα was assessed by ELISA to determine the half-maximal effective concentration (EC_50_) of each antibody format: IgG (Adalimumab), monomeric ABC-301 and oligomeric ABC-301. The measured EC_50_ values were 0.11 ± 0.02 µg/mL for Adalimumab, 0.17 ± 0.03 µg/mL for monomeric ABC-301 and 0.18 ± 0.06 µg/mL for oligomeric ABC-301 (FIG 2A.). These comparable EC_50_ values demonstrate that binding to human TNFα is preserved following reformatting from the IgG to the sIgA1 isotype, supporting the ability of the platform to maintain the antigen-binding properties of the parental antibody throughout the transformation process.

Functional TNFα neutralization was evaluated using TNFα-sensitive L929 fibroblast-like cells. TNFα induces cell death in this cell line, whereas the presence of anti-TNFα antibodies inhibits TNFα-mediated apoptosis, thereby providing a functional measure of the neutralizing activity of each antibody format. ABC-301 monomer and oligomer fractions were compared with Adalimumab. The half-maximal inhibitory concentration (IC_50_) was 0.14 ± 0.02 µg/mL for Adalimumab, 0.10 ± 0.01 µg/mL for monomeric ABC-301 and 0.13 ± 0.02 µg/mL for oligomeric ABC-301 (Fig. 2B). Given that the molecular weight of an IgG is approximately 150kDa and that of monomeric sIgA is approximately 350kDa, the IC_50_ values correspond to approximately 0.93 nM for Adalimumab and 0.29 nM for monomeric ABC-301. On a molar basis, ABC-301 therefore exhibits an approximately threefold greater TNFα-neutralizing potency than its parental IgG format, consistent with the increased valency conferred by the dimeric, tetravalent architecture of sIgA.

**Figure 2.**
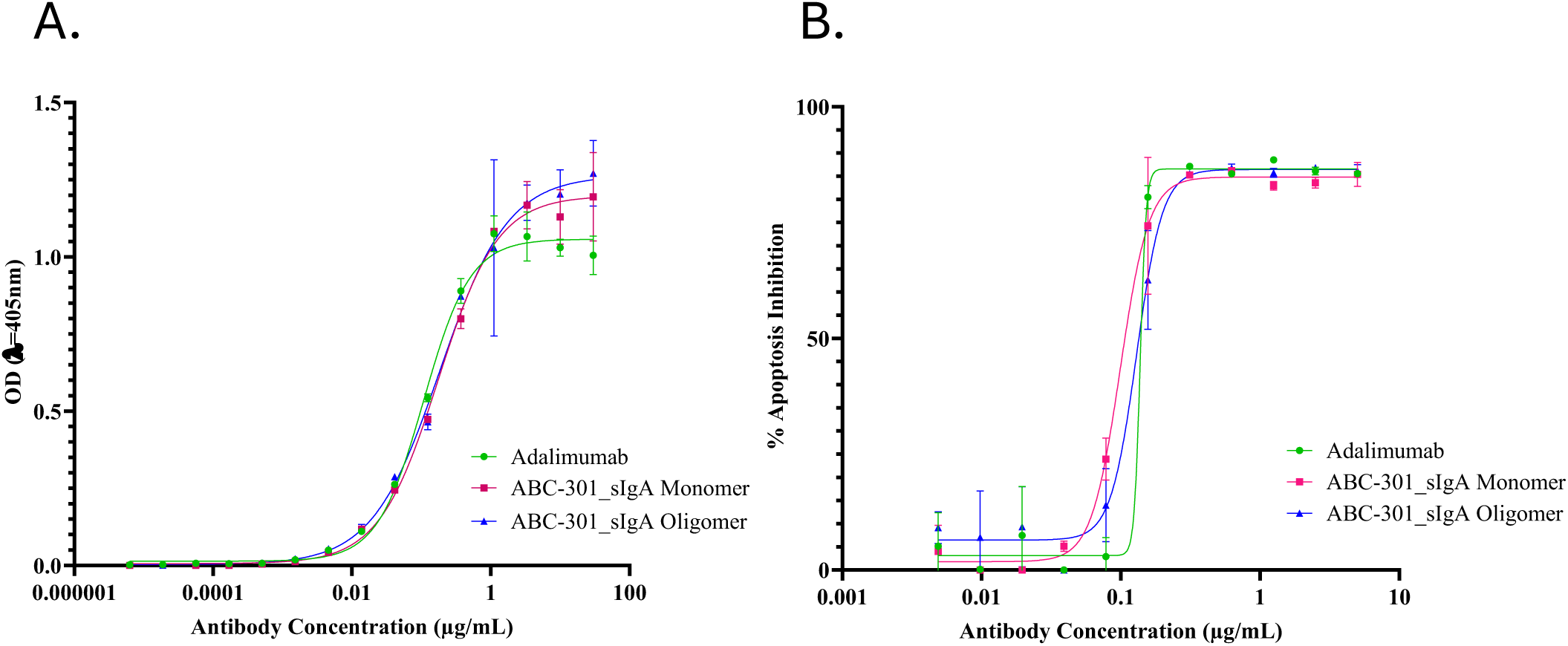
Preservation of human TNFα binding and neutralizing activity following IgG-to-sIgA reformatting. **(A)** Concentration-dependent binding of Adalimumab and ABC-301 preparations to recombinant human tumor necrosis factor-alpha (TNFα), determined by enzyme-linked immunosorbent assay (ELISA). The predominant dimeric sIgA-enriched fraction and oligomer-enriched ABC-301 fraction were compared with parental Adalimumab IgG1. Half-maximal effective concentrations (EC_50_) were 0.11 ± 0.02 µg/mL for Adalimumab, 0.17 ± 0.03 µg/mL for the predominant ABC-301 fraction, and 0.18 ± 0.06 µg/mL for the oligomer-enriched fraction. **(B)** Functional neutralization of human TNFα in the L929 cell-based cytotoxicity assay. TNFα-induced cell death was assessed in the presence of increasing concentrations of each antibody preparation, and half-maximal inhibitory concentrations (IC_50_) were determined from concentration-response curves. IC_50_ values were 0.14 ± 0.02 µg/mL for Adalimumab, 0.10 ± 0.01 µg/mL for the predominant ABC-301 fraction, and 0.13 ± 0.02 µg/mL for the oligomer-enriched fraction. Based on approximate molecular masses of 150 kDa for Adalimumab and 350 kDa for sIgA ABC-301, the corresponding whole-molecule IC_50_ values were approximately 0.93 and 0.29 nM, respectively.

### Orally administered ABC-301 is concentrated in the gastrointestinal tract

After a single oral dose in healthy mice, ABC-301-associated radioactivity was detected predominantly in gastrointestinal compartments. In biodistribution experiment, the highest early signals were observed in stomach and small intestine, with substantial recovery in gastrointestinal washes. At 2 h, stomach and small-intestinal levels were 26.0 ± 8.1% injected dose per gram and 4.0 ± 1.3% injected dose per gram, respectively, while gastrointestinal washes contained 36.5 ± 5.5% of the injected dose. Blood contained 7.9 ± 0.5% of the injected dose at 2 h, with lower early signals in liver and spleen. Radioactivity progressively declined after 2 h and was minimal in most compartments by 48 h. In healthy-mouse pharmacokinetic measurements, systemic exposure peaked within the first hours after dosing and declined markedly by 24 h. DSS-treated animals showed higher and more prolonged systemic exposure than healthy animals, consistent with disruption of the intestinal barrier, but gastrointestinal localization remained a dominant feature of the distribution profile.

### Oral ABC-301 attenuates DSS-induced colon shortening

ABC-301 was well tolerated over the seven-day oral dosing period at all tested doses (2, 1, 0.5 and 0.2 mg/kg) (Fig. 3A). To induce the inflammatory and physiological features of ulcerative colitis, including colon shortening, DSS was administered in the drinking water for five days and then withdrawn for the entire seven-day treatment period. DSS exposure induced body-weight loss beginning around days 5–8, followed by spontaneous stabilization and recovery in untreated animals (grey curve, Fig. 3B). The magnitude of body-weight loss was markedly reduced in animals treated daily with oral ABC-301, with body weight stabilizing as early as day 8 and fully recovering by day 13, at the endpoint. Animals receiving Adalimumab as a single 2 mg/kg subcutaneous dose on day 5 also showed less pronounced body-weight loss than untreated animals, although the loss remained greater than that observed with oral ABC-301. From day 9 onwards, however, the Adalimumab-treated animals followed a body weight recovery trajectory comparable to that of the ABC-301-treated group.

**Figure 3.**
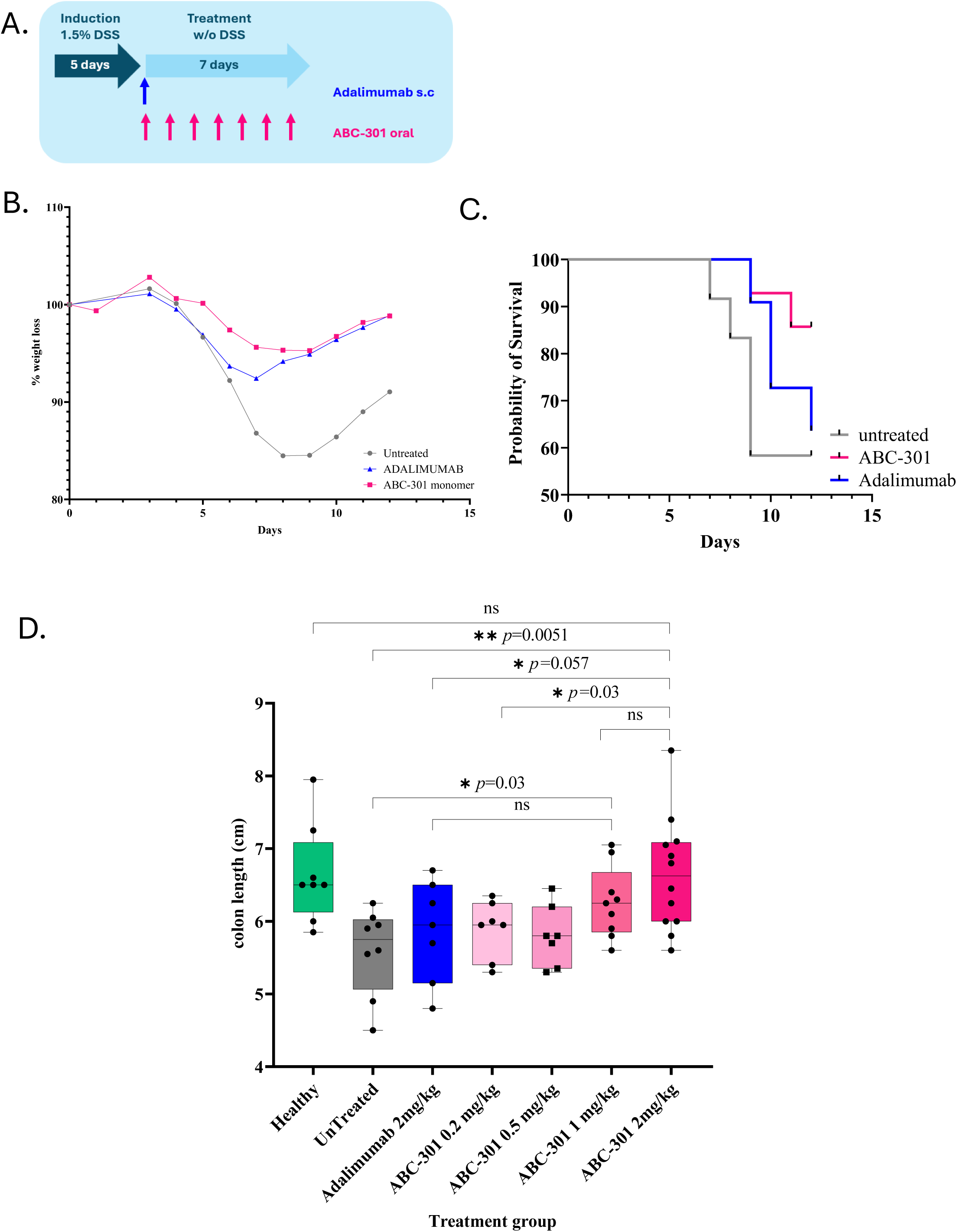
Experimental design and effects of orally administered ABC-301 on clinical and anatomical outcomes in DSS-induced colitis. **(A)** Schematic representation of the experimental protocol. Colitis was induced in C57BL/6 mice transgenic for human FcαRI (CD89) by administration of 1.5% dextran sulfate sodium (DSS) in drinking water for five consecutive days. Following DSS withdrawal, ABC-301 was administered by oral gavage once daily for seven days (days 6–12), whereas Adalimumab was administered as a single subcutaneous injection at 2 mg/kg on day 6. Untreated DSS-exposed mice and healthy animals served as controls. **(B)** Longitudinal changes in body weight in DSS-exposed mice receiving oral ABC-301 (2mg/kg/day), subcutaneous Adalimumab (2mg/kg, single dose), or no antibody treatment. Body weight was monitored throughout DSS induction and the subsequent treatment period. **(C)** Kaplan–Meier survival curves showing the proportion of surviving animals during the experimental period in the indicated treatment groups. **(D)** Colon length measured at the experimental endpoint (day 13) in healthy mice, untreated DSS-exposed mice, and DSS-exposed mice receiving Adalimumab or ABC-301 at the indicated doses.

Mortality in the untreated group began around day 7 and increased rapidly, resulting in approximately 60% survival by day 9, which remained unchanged until the endpoint of the study (FIG 3C.). In contrast, ABC-301-treated animals maintained 100% survival until approximately day 9 and showed approximately 85% survival at day 13. Adalimumab treatment provided an intermediate survival profile, with survival declining from day 9 onwards to approximately 65% by the end of the experiment. Daily oral ABC-301 treatment seems associated with improved survival and delayed mortality compared with untreated animals and Adalimumab treatment.

In the DSS-induced mouse model of ulcerative colitis, colon length is an indicator of intestinal inflammation severity. In healthy animals, median colon length was approximately 6.5 ± 0.7 cm, whereas DSS exposure resulted in marked colon shortening in untreated animals, with a colon length of 5.7 ± 0.6 cm, corresponding to a 12.5% reduction. At 2 mg/kg, oral ABC-301 significantly attenuated DSS-induced colon shortening compared with untreated DSS controls (*p* = 0.0068), restoring colon length to values comparable to those of healthy animals (6.6 ± 0.8 cm vs 6.5 ± 0.7 cm, respectively; *p* = 0.909). In contrast, animals treated with Adalimumab at 2 mg/kg exhibited a 10% reduction in colon length compared with the oral ABC-301-treated group. Although this difference suggested a greater anti-inflammatory effect of ABC-301, it did not reach conventional statistical significance (*p* = 0.057). Finally, in the dose-ranging experiment, 2 and 1 mg/kg oral ABC-301 produced the clearest preservation of colon length, whereas 0.5 and 0.2 mg/kg produced smaller effects.

### Oral ABC-301 Demonstrates Anti-Inflammatory Activity

Monocyte recruitment, which contributes to tissue macrophage replenishment, and the accumulation of pro-inflammatory M1 macrophages are key cellular markers of intestinal inflammation. These myeloid cell populations were therefore investigated in colonic tissues to assess the anti-inflammatory activity of ABC-301 (Fig 4). In untreated DSS-exposed animals, the proportions of monocytes (Fig 4C.), total macrophages (Fig 4A.) and pro-inflammatory M1 macrophages (Fig 4B.) were markedly increased compared with healthy animals (1.5% [1.7] vs 0.1% [0.2], *p*=0.0023; 1.1% [1.2] vs 0.07% [0.09], *p*=0.0025; and 1.1% [1.1] vs 0.06% [0.09], *p*=0.0003, respectively).

**Figure 4.**
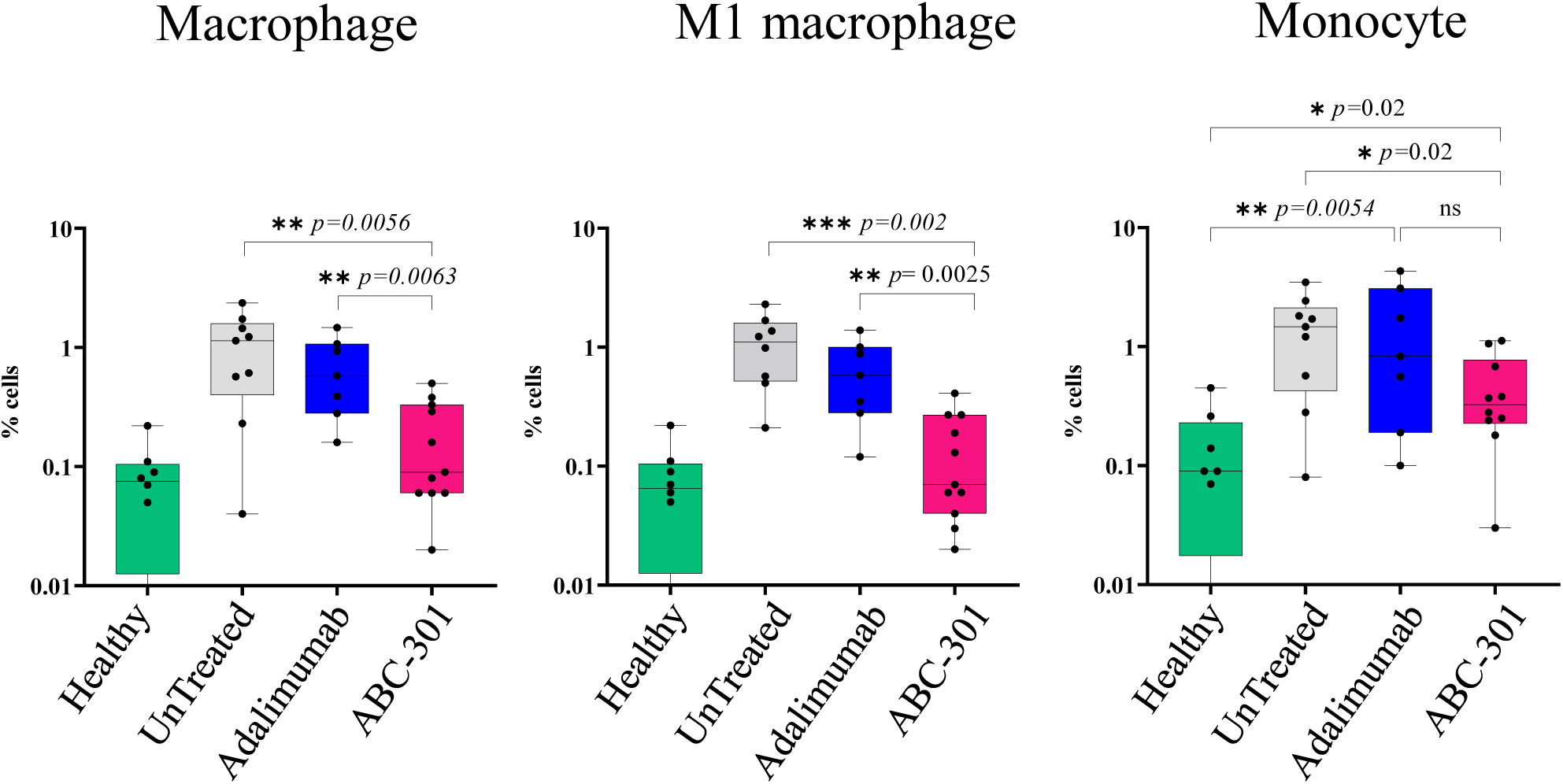
Oral ABC-301 treatment is associated with reduced accumulation of inflammatory myeloid-cell populations in colonic tissue. Flow-cytometric quantification of myeloid-cell populations isolated from colonic tissues at day 13 following DSS-induced colitis and treatment with oral ABC-301 or subcutaneous Adalimumab. **(A)** Total macrophages, identified according to CD11c, CD11b, F4/80, and Ly6G gating strategy. **(B)** CD80-positive inflammatory macrophages, designated as M1-like macrophages, identified within the CD11c^+^CD11b^+^F4/80^+^ population. **(C)** Monocytes, identified as CD11b^+^Ly6G^−^F4/80^−^ cells. Cell populations were quantified after exclusion of debris, doublets, and nonviable cells and expressed as percentages of the specified viable-cell reference population.

Oral ABC-301 treatment significantly reduced the accumulation of these inflammatory myeloid populations compared with untreated group, resulting in an 80% reduction in monocytes (*p*=0.02), a 92% reduction in total macrophages (*p*=0.0056) and a 94% reduction in pro-inflammatory M1 macrophages (*p*=0.002). Notably, ABC-301 induced a significantly greater reduction in colonic M1 macrophages than its parental IgG counterpart, Adalimumab (*p*=0.0025), demonstrating superior suppression of this key pro-inflammatory cell population following oral administration.

### ABC-301 reduces tissue calprotectin and lipocalin-2

DSS-induced colitis resulted in increased levels of both calprotectin and lipocalin-2 compared with healthy controls, in both colonic tissue (Fig. 5A,C) and fecal samples (Fig. 5B,D). In colonic tissue, oral ABC-301 treatment significantly reduced calprotectin levels by approximately 45-fold compared with untreated controls (49.7 ng/g [676.4 ng/g] vs 2.2 µg/g [1.2 µg/g], respectively; *p* = 0.0048), restoring concentrations to levels comparable to those observed in healthy animals (13.2 ng/g [27.5 ng/g]; not statistically significant vs healthy controls). In contrast, Adalimumab treatment failed to normalize tissue calprotectin levels, which remained approximately 22-fold higher than those measured in healthy animals (289.7 ng/g [483.0 ng/g] vs 13.2 ng/g [27.5 ng/g], respectively; *p* = 0.0003).

**Figure 5.**
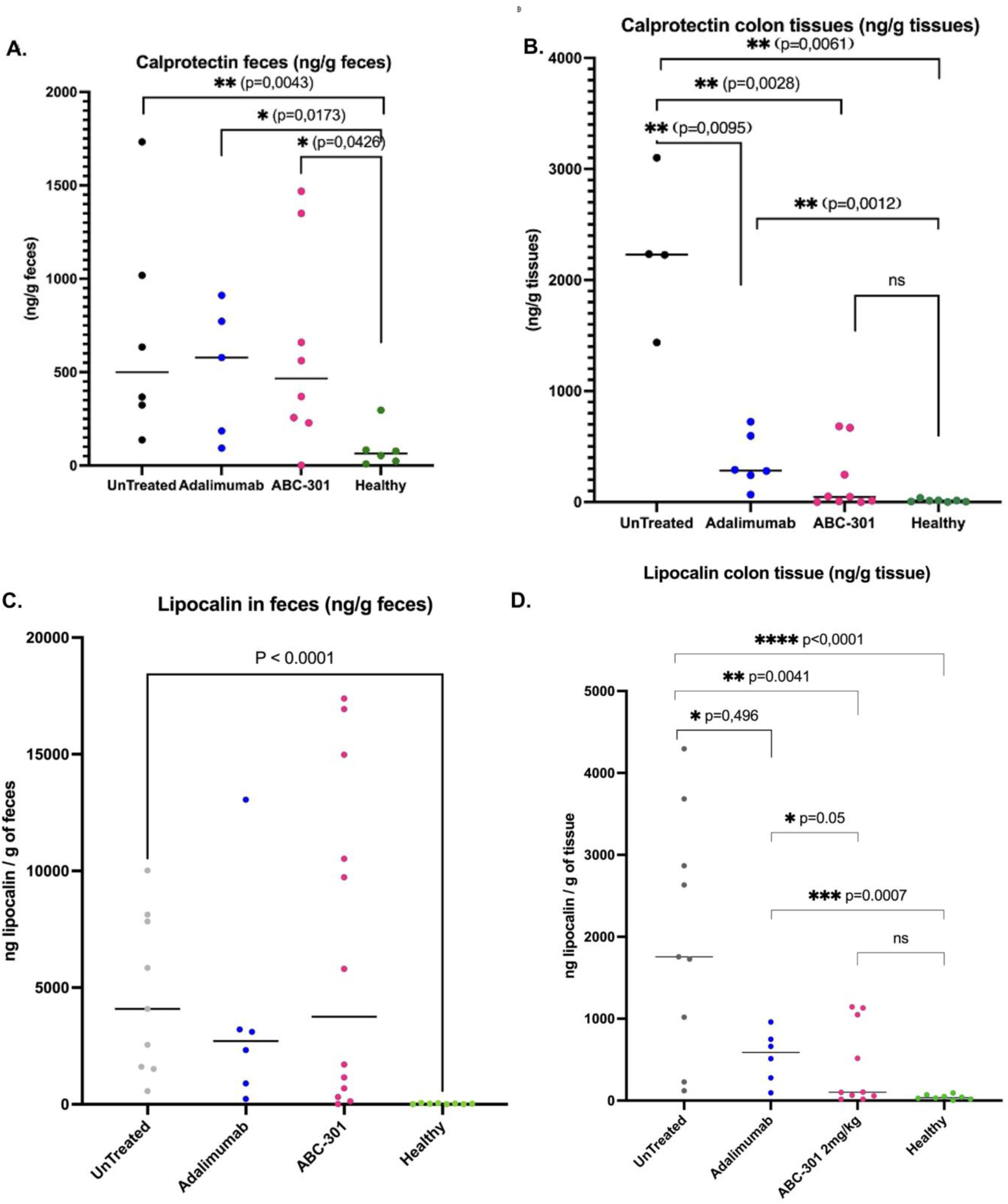
Effects of oral ABC-301 on calprotectin and lipocalin-2 concentrations in fecal samples and intestinal tissue during DSS-induced colitis. Quantification of inflammatory biomarkers in fecal samples and colonic tissue collected from healthy mice, untreated DSS-exposed mice, and DSS-exposed mice treated with oral ABC-301 or subcutaneous Adalimumab. **(A)** Fecal calprotectin concentrations **(B)** Intestinal-tissue calprotectin concentrations **(C)** Fecal lipocalin-2 concentrations and **(D)** Intestinal-tissue lipocalin-2 concentrations measured by ELISA in feces or in colonic tissue extracts.

**Figure 6.**
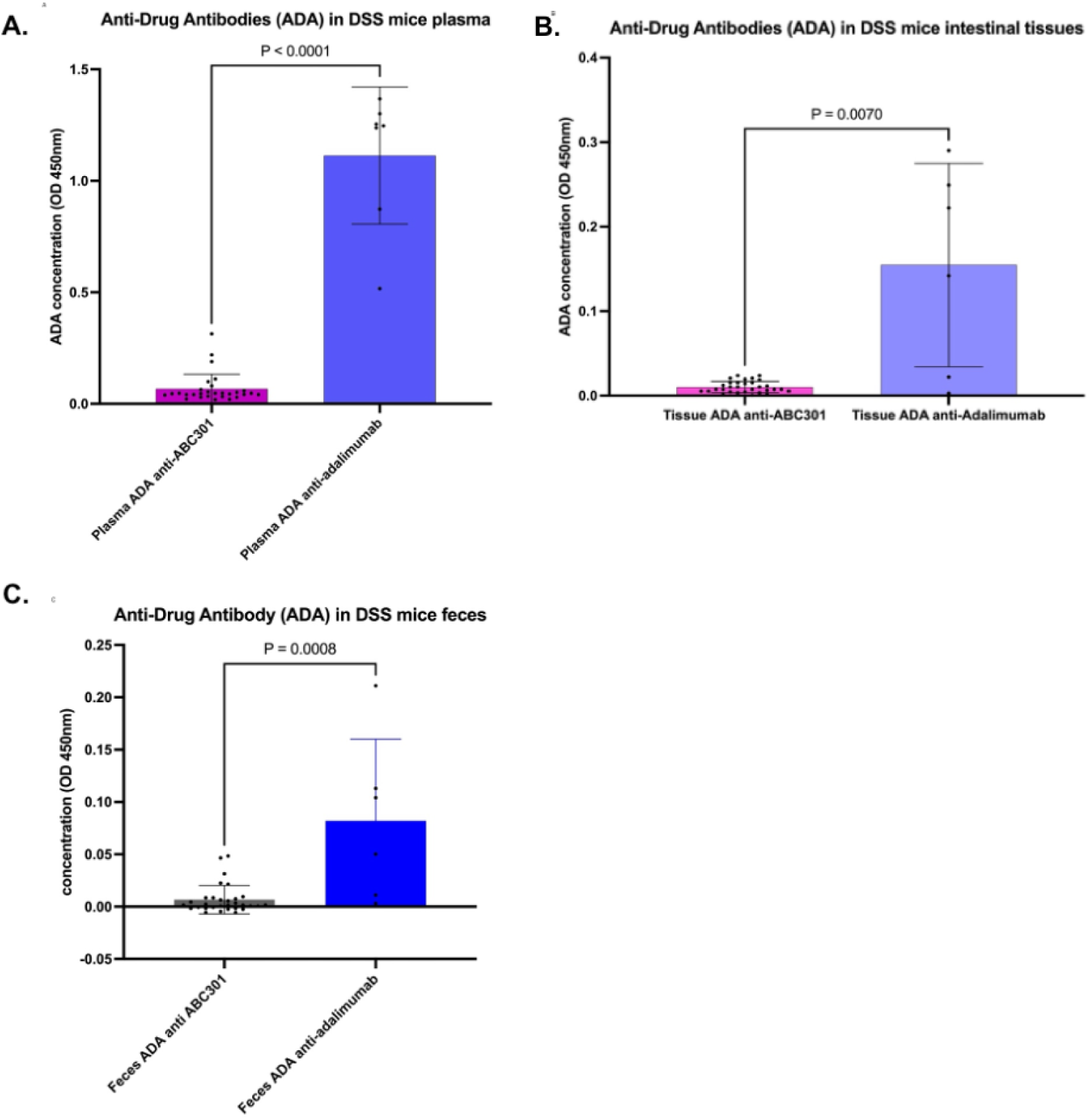
Exploratory assessment of anti-drug antibody responses following oral ABC-301 or subcutaneous Adalimumab administration. Anti-drug antibody (ADA) measurements in biological samples collected from DSS-exposed human CD89-transgenic mice following seven consecutive daily oral doses of ABC-301 (0.2, 0.5, 1, or 2 mg/kg/day) or a single subcutaneous injection of Adalimumab (2mg/kg). ADA detection in plasma (A), in intestinal-tissue preparations (B) and in fecal samples (C). ABC-301-specific and Adalimumab-specific antibody responses were assessed using their respective antigen-dependent ELISA assays.

A similar pattern was observed for lipocalin-2. Oral ABC-301 treatment restored colonic tissue lipocalin-2 levels to values comparable to those observed in healthy animals (102.0 ng/g [1.0 µg/g] vs 35.3 ng/g [46.3 ng/g], respectively; not statistically significant). In contrast, following subcutaneous Adalimumab treatment, tissue lipocalin-2 concentrations remained approximately 17-fold higher than those in healthy controls (587.0 ng/g [570.7 ng/g] vs 35.3 ng/g [46.3 ng/g], respectively; *p* = 0.0007) and approximately sixfold higher than those observed in ABC-301-treated animals.

Finally, neither ABC-301 nor Adalimumab measurably reduced fecal calprotectin or lipocalin-2 levels. This dissociation between tissue and fecal inflammatory biomarkers may reflect delayed luminal clearance relative to the resolution of local tissue inflammation.

### Oral ABC-301 induces a significantly lower Anti-Drug Antibody response than subcutaneous Adalimumab

Murine anti-ABC-301 antibodies (Anti-Drug Antibodies, ADAs) were quantified using an ABC-301-specific ELISA in C57BL/6 transgenic hCD89 mice treated orally with ABC-301 at daily doses of 2, 1, 0.5 or 0.2 mg/kg for seven consecutive days. Similarly, anti-Adalimumab antibodies were measured in animals receiving a single subcutaneous injection of Adalimumab (2 mg/kg).

When data from all ABC-301 dose groups were pooled, ADA levels were significantly lower in ABC-301-treated animals than in Adalimumab-treated animals across all three compartments analyzed: plasma (*p* < 0.0001) (FIG 6A.), intestinal tissue (*p*=0.0070) (FIG 6B.) and feces (*p*=0.0008) (FIG 6C.). No clear dose-dependent increase in ADA levels was observed across the tested ABC-301 doses. Furthermore, circulating anti-ABC-301 antibody levels were not associated with correspondingly elevated ADA levels in intestinal tissue, the primary site of ABC-301 therapeutic activity.

Together, a significantly lower Anti-Drug Antibody response following repeated oral administration of ABC-301 is demonstrated in comparison with subcutaneous Adalimumab single-dose administration in immunocompetent C57BL/6 transgenic hCD89 mice, supporting a favorable relative immunogenicity profile for the oral sIgA format.

### Reduced susceptibility of ABC-301 to functional neutralization by patient-derived anti-Adalimumab antibodies

The functional consequences of anti-adalimumab antibodies from seven treated patients were evaluated using paired comparisons of adalimumab and ABC-301 in an L929 TNFa-dependent apoptosis assay. At the threshold at which ADA-mediated neutralization was no longer detectable, the molar ratios of therapeutic antibody (Adalimumab or ABC-301) to ADA were determined and are presented in Figure 7. It required significantly higher therapeutic antibody-to-ADA ratios to overcome ADA-mediated inhibition of Adalimumab than of ABC-301. Ratio paired t-test analysis revealed a 17.5-fold difference in the geometric mean antibody-to-ADA ratio (95% CI: 3.30–92.88; p = 0.0057) from 2-fold difference for the patient #5 to 150-fold difference for the patient #1, with consistently lower ratios observed for ABC-301 across all seven patients. For example, ADAs present in plasma samples from patients #1, #2 and #3 exhibited rather no inhibitory activity against ABC-301, with no detectable ADA-mediated neutralization at molar ratios of 0.91, 1.1 and 2.2, respectively. In contrast, Adalimumab was, on average, 104-fold more susceptible to ADA-mediated neutralization, with corresponding ADA-to-Adalimumab molar ratios of 137.2, 32.0 and 142.9, respectively. The nature of the ADAs generated appears to vary substantially between patients, revealing distinct neutralization profiles: (i) negligible neutralization of ABC-301 but strong neutralization of Adalimumab (patients #1, #2 and #3); (ii) intermediate neutralization of ABC-301 and Adalimumab (patients #4 and #5); (iii) strong neutralization of ABC-301 and very strong neutralization of Adalimumab (patient #6); and (iv) similarly pronounced neutralization of both ABC-301 and Adalimumab (patient #7). Interpatient heterogeneity in ADA-mediated functional inhibition potentially reflects differences in antibody specificity, epitope recognition, and neutralizing capacity. However, despite this interpatient heterogeneity, these findings demonstrate that reformatting the Adalimumab isotype into an sIgA1 architecture globally reduces its susceptibility to functional inhibition by anti-Adalimumab antibodies in this *in vitro* assay.

**Figure 7.**
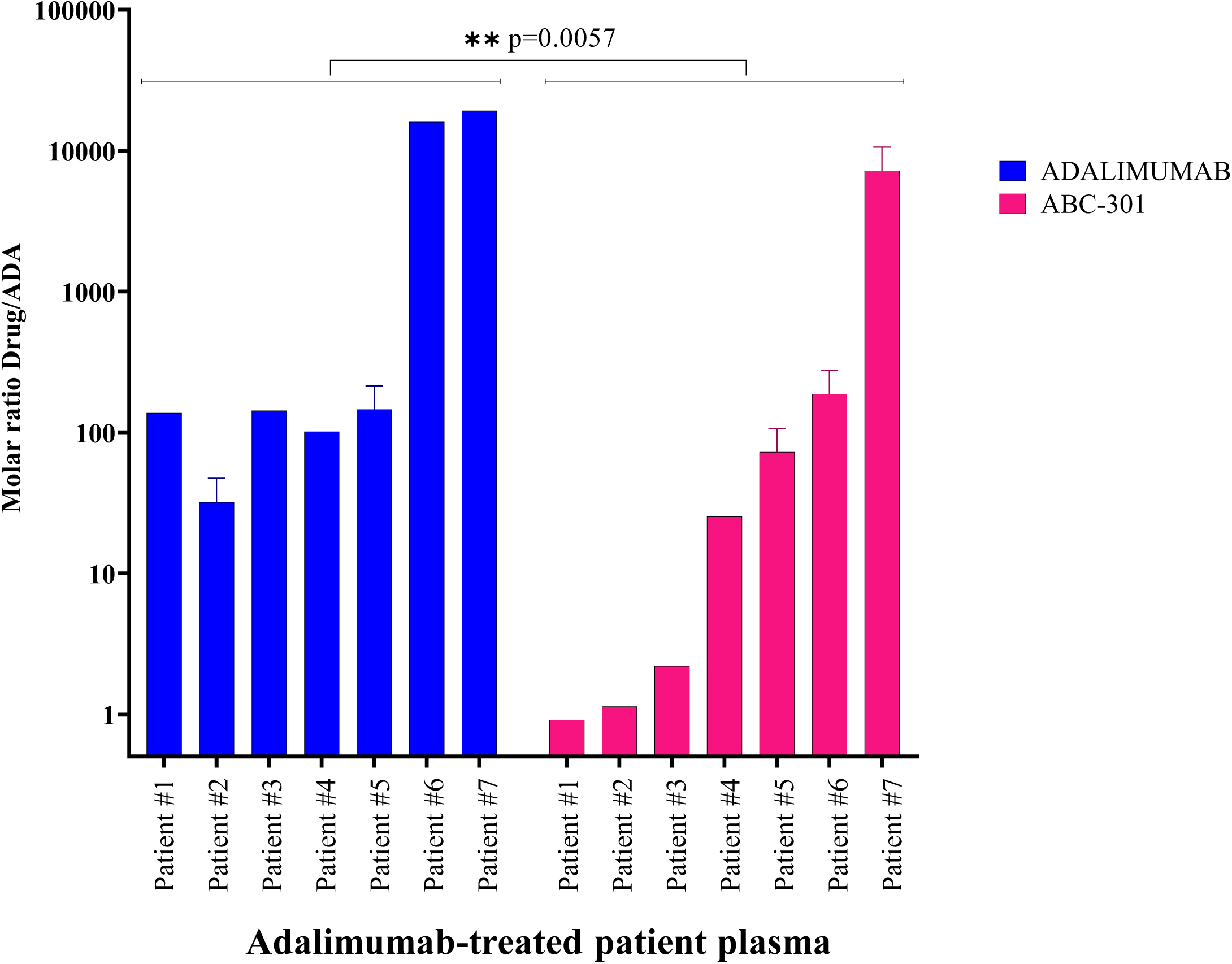
Comparative susceptibility of ABC-301 and Adalimumab to neutralization by patient-derived anti-Adalimumab antibodies. The neutralizing activity of anti-drug antibodies (ADAs) obtained from seven patients previously treated with Adalimumab was evaluated against parental Adalimumab IgG1 and its secretory IgA counterpart, ABC-301, using a TNFα-dependent L929 cell cytotoxicity assay. Both therapeutic antibodies were tested at an identical molar concentration (2 × 10⁻¹⁰ M) in the presence of increasing concentrations of patient-derived ADAs and recombinant human TNFα (7.5 ng/mL). ADA-mediated inhibition of therapeutic antibody activity was assessed by measuring the restoration of TNFα-induced cytotoxicity. For each patient, the ADA concentration required to inhibit the protective activity of Adalimumab or ABC-301 was determined from the corresponding concentration-response curves. The relative susceptibility of the two antibodies to ADA-mediated neutralization was expressed as the ratio of ADA concentrations required to inhibit Adalimumab versus ABC-301.

## DISCUSSION

The present study provides preclinical proof of concept for converting a clinically validated therapeutic IgG into an orally active secretory IgA (sIgA). By incorporating the Adalimumab-derived anti-TNFα paratope into an sIgA1 backbone, ABC-301 preserved antigen binding and TNFα neutralization *in vitro*, while demonstrating anti-inflammatory activity following oral administration in DSS-induced colitis. These findings establish an experimental foundation for an adaptable antibody-engineering platform designed to modify therapeutic antibody architecture, delivery, and tissue distribution. Despite substantial differences in molecular organization, valency, and Fc structure, ABC-301 retained the functional properties of its parental IgG. On a molar basis, ABC-301 exhibited approximately threefold greater TNFα-neutralizing potency than Adalimumab, consistent with a potential contribution of increased sIgA valency. Preservation of antigen recognition and biological activity following reformatting represents a prerequisite for extending this strategy to other clinically validated therapeutic antibodies.

Oral ABC-301 produced substantial improvements in critical biomarkers of intestinal inflammation, including colon length, inflammatory myeloid-cell accumulation, and tissue concentrations of calprotectin and lipocalin-2. At 2 mg/kg, ABC-301 significantly attenuated DSS-induced colon shortening, restoring colon length to values comparable to healthy controls. Relative to the DSS-induced deficit, ABC-301 achieved 117% recovery, compared with 27% following subcutaneous Adalimumab. Although this between-treatment difference did not reach statistical significance (*p* = 0.057), the findings suggest that ABC-301 has a strong potential to preserve intestinal integrity. These anatomical improvements were accompanied by reduced colonic monocyte and macrophage accumulation, particularly pro-inflammatory M1 macrophages, whose abundance was significantly lower following ABC-301 treatment than with Adalimumab. Colonic calprotectin and lipocalin-2 concentrations also decreased towards healthy values, supporting attenuation of local innate inflammation. However, corresponding reductions were not observed in fecal biomarkers during the study period, potentially reflecting different kinetics of tissue recovery and luminal protein clearance. Longitudinal investigations will be required to clarify this discrepancy.

Biodistribution analysis further supports the intended mucosa-directed pharmacology. Following oral administration, the highest early ^125^I-associated signals were detected in gastrointestinal compartments, whereas systemic radioactivity remained comparatively limited and transient, including under inflammatory conditions. This profile suggests preferential gastrointestinal exposure, although radiolabel detection does not establish the presence of intact, biologically active antibody. Future studies should quantify molecular integrity and local TNFα target engagement.

Repeated oral ABC-301 administration also induced significantly lower anti-drug antibody (ADA) responses than subcutaneous Adalimumab in mice, across plasma, intestinal tissue and feces. Although these findings are consistent with the tolerogenic properties of the intestinal mucosa, differences in antibody format, administration route, dosing schedule and species-specific immune recognition must be carefully considered before definitively concluding that orally administered sIgA exhibits lower immunogenicity.

Importantly, ABC-301 exhibited significantly reduced susceptibility to functional inhibition by anti-Adalimumab antibodies from Adalimumab-treated patients, despite sharing the same paratope. This observation suggests that ADA-mediated inhibition is not determined exclusively by recognition of the shared antigen-binding region but may also depend on epitopes or structural features modified by IgG-to-sIgA1 reformatting. Recognition of IgG constant-region determinants remains a plausible mechanism, potentially promoting hepatic clearance of Adalimumab–ADA immune complexes and thereby contributing to the loss of therapeutic efficacy. These findings raise the possibility that ABC-301 could retain activity in patients experiencing ADA-associated loss of response to Adalimumab. Furthermore, oral delivery may potentially limit treatment-induced immunogenicity and support prolonged administration in chronic inflammatory diseases; both possibilities require further clinical validation.

Beyond ABC-301, the principal technological significance of this study lies in demonstrating that a clinically validated IgG paratope can be transferred into a secretory IgA architecture while preserving biological functionality. This strategy could enable selected therapeutic antibodies to exploit sIgA-specific properties, including multivalency, mucosal stability, and enhanced tissue distribution. Potential applications extend to inflammatory, infectious, immune-mediated and neoplastic diseases, particularly when local target engagement offers advantages over sustained systemic exposure. However, IgG-to-sIgA conversion should not be considered universally applicable. Each candidate requires optimization according to its antigen, mechanism of action, and intended indication. Target accessibility and binding properties are critical, although therapeutic activity need not necessarily remain confined to mucosal compartments, as previous studies detected orally administered anti-CEACAM5 sIgA in pulmonary and hepatic metastases in *in vivo* human CRC model. Molecular geometry, valency, glycosylation, oligomerization and aggregation propensity may also influence functional activity, stability, manufacturability, and tissue exposure.

Particular attention must be given to FcαRI (CD89) engagement. Unlike IgG, IgA can interact with FcαRI-expressing neutrophils, monocytes and macrophages, potentially triggering cytotoxicity, phagocytosis, or inflammatory mediator release. Such responses may be therapeutically beneficial, as previously suggested for anti-CEACAM5 sIgA in colorectal cancer, or undesirable in inflammatory diseases. Although ABC-301 administration in human CD89-transgenic mice was not associated with detectable hyperactivation of CD89-positive neutrophils, candidate-specific evaluation of immune-complex formation, FcαRI signaling and tissue toxicity remains essential.

Several limitations warrant consideration. DSS-induced colitis primarily models acute epithelial injury and innate inflammation rather than the chronic immunopathology of human inflammatory bowel disease. Confirmation in complementary chronic or immune-mediated models is needed. Exposure-matched comparisons, appropriate administration-route controls, and validation of TNFα target engagement in the experimental system are also required to distinguish format-dependent effects from differences in pharmacological exposure. Long-term studies should further assess immunogenicity, FcαRI-dependent inflammation, and systemic safety.

Collectively, these findings establish ABC-301 as a preclinical demonstration of an adaptable IgG-to-sIgA engineering strategy. By combining preservation of a clinically validated paratope with oral delivery, mucosa-directed activity and reduced susceptibility to pre-existing anti-Adalimumab antibodies, this approach may expand the therapeutic potential of selected IgGs whose clinical application is constrained by poor tissue accessibility or toxic prolonged systemic exposure.

## DECLARATIONS

Ethics statement. Animal procedures were conducted in accordance with Directive 2010/63/EU and French Decree 2013-118. The dose-ranging study protocol was reviewed and approved by the local ethics committee under APAFIS application APAFIS #56954-2025091016282834 v3_17/11/2025. Animals were randomly allocated to different experimental groups. Investigators performing histological, imaging, and flow cytometric analyses were blinded to treatment allocation whenever feasible. No animals were excluded from the analyses unless predefined experimental criteria were met.

## ACKNOWLEDGEMENTS

We would like to thank Dr Eric Pinaud for making the research facilities at the CRIBL laboratory available to us. We are grateful to BISCEm unit (Univ. Limoges, UAR 2015 CNRS, US 42 Inserm, CHU Limoges) for technical support regarding cytometry experiments and animal core facility. We would like to thank Professor Stéphane Paul for helpful discussions and for providing plasma samples from Adalimumab-treated patients and for quantifying anti-drug antibodies (ADAs) in these samples.

## CONTRIBUTORS

Conceptualization: AC and JMH, Methodology: AC. Formal Analysis: AC, MB, JMH. Investigation: MB, AP, AC. Resources: JMH and AC. Writing Original Draft: AC and JMH. Writing, Review and Editing: BL, JMH, AC. Supervision and Funding Acquisition: JMH and AC. Guarantor: JMH and AC. JMH and AC contributed equally to this work.

## REFERENCES

1. Lockhart-Mummery HE, Morson BC. Crohn’s disease (regional enteritis) of the large intestine and its distinction from ulcerative colitis. Gut. 1960 Jun;1(2):87–105. doi:10.1136/gut.1.2.87 PubMed PMID: 14417801; PubMed Central PMCID: PMC1413217.

2. Neurath MF. Strategies for targeting cytokines in inflammatory bowel disease. Nat Rev Immunol. 2024 Aug;24(8):559–76. doi:10.1038/s41577-024-01008-6

3. Braegger CP, Nicholls S, Murch SH, Stephens S, MacDonald TT. Tumour necrosis factor alpha in stool as a marker of intestinal inflammation. Lancet. 1992 Jan 11;339(8785):89–91. doi:10.1016/0140-6736(92)90999-j PubMed PMID: 1345871.

4. van Dullemen HM, van Deventer SJ, Hommes DW, Bijl HA, Jansen J, Tytgat GN, et al. Treatment of Crohn’s disease with anti-tumor necrosis factor chimeric monoclonal antibody (cA2). Gastroenterology. 1995 Jul;109(1):129–35. doi:10.1016/0016-5085(95)90277-5 PubMed PMID: 7797011.

5. Targan SR, Hanauer SB, van Deventer SJ, Mayer L, Present DH, Braakman T, et al. A short-term study of chimeric monoclonal antibody cA2 to tumor necrosis factor alpha for Crohn’s disease. Crohn’s Disease cA2 Study Group. N Engl J Med. 1997 Oct 9;337(15):1029–35. doi:10.1056/NEJM199710093371502 PubMed PMID: 9321530.

6. Ungar B, Engel T, Yablecovitch D, Lahat A, Lang A, Avidan B, et al. Prospective Observational Evaluation of Time-Dependency of Adalimumab Immunogenicity and Drug Concentrations: The POETIC Study. Am J Gastroenterol. 2018 Jun;113(6):890–8. doi:10.1038/s41395-018-0073-0 PubMed PMID: 29867175.

7. Vande Casteele N, Herfarth H, Katz J, Falck-Ytter Y, Singh S. American Gastroenterological Association Institute Technical Review on the Role of Therapeutic Drug Monitoring in the Management of Inflammatory Bowel Diseases. Gastroenterology. 2017 Sep;153(3):835–857.e6. doi:10.1053/j.gastro.2017.07.031 PubMed PMID: 28774547.

8. Yanai H, Lichtenstein L, Assa A, Mazor Y, Weiss B, Levine A, et al. Levels of drug and antidrug antibodies are associated with outcome of interventions after loss of response to infliximab or adalimumab. Clin Gastroenterol Hepatol. 2015 Mar;13(3):522–530.e2. doi:10.1016/j.cgh.2014.07.029 PubMed PMID: 25066837.

9. Deyhim T, Cheifetz AS, Papamichael K. Drug Clearance in Patients with Inflammatory Bowel Disease Treated with Biologics. J Clin Med. 2023 Nov 16;12(22):7132. doi:10.3390/jcm12227132 PubMed PMID: 38002743; PubMed Central PMCID: PMC10672599.

10. Yarur AJ, Jain A, Sussman DA, Barkin JS, Quintero MA, Princen F, et al. The association of tissue anti-TNF drug levels with serological and endoscopic disease activity in inflammatory bowel disease: the ATLAS study. Gut. 2016 Feb;65(2):249–55. doi:10.1136/gutjnl-2014-308099 PubMed PMID: 25670812.

11. Kennedy NA, Heap GA, Green HD, Hamilton B, Bewshea C, Walker GJ, et al. Predictors of anti-TNF treatment failure in anti-TNF-naive patients with active luminal Crohn’s disease: a prospective, multicentre, cohort study. Lancet Gastroenterol Hepatol. 2019 May;4(5):341–53. doi:10.1016/S2468-1253(19)30012-3 PubMed PMID: 30824404.

12. Harnik S, Abitbol CM, Haj Natour O, Yavzori M, Fudim E, Picard O, et al. Prospective Observational Evaluation of the Time-Dependency of Adalimumab Immunogenicity and Drug Concentration in Ulcerative Colitis Patients: the POETIC II Study. J Crohns Colitis. 2024 Mar 1;18(3):341–8. doi:10.1093/ecco-jcc/jjad156 PubMed PMID: 37691574.

13. Tournier Q, Paul S, Williet N, Berger AE, Veyrard P, Boschetti G, et al. Early detection of anti-drug antibodies during initiation of anti-tumour necrosis factor therapy predicts treatment discontinuation in inflammatory bowel disease. Aliment Pharmacol Ther. 2021 Jun;53(11):1190–200. doi:10.1111/apt.16333 PubMed PMID: 33872404.

